# Relative time constraints improve molecular dating

**DOI:** 10.1101/2020.10.17.343889

**Authors:** Gergely J. Szöllősi, Sebastian Höhna, Tom A. Williams, Dominik Schrempf, Vincent Daubin, Bastien Boussau

## Abstract

Dating the tree of life is central to understanding the evolution of life on Earth. Molecular clocks calibrated with fossils represent the state of the art for inferring the ages of major groups. Yet, other information on the timing of species diversification can be used to date the tree of life. For example, horizontal gene transfer events and ancient coevolutionary relationships such as (endo)symbioses can imply temporal relationships between two nodes in a phylogeny (Davín et al. 2018). These alternative sources of temporal constraints can be particularly helpful when the geological record is sparse, e.g. for microorganisms, which represent the vast majority of extant and extinct biodiversity.

Here, we present a new method to combine fossil calibrations and relative age constraints to estimate chronograms. We provide an implementation of relative age constraints in RevBayes (Höhna et al. 2016) that can be combined in a modular manner with the wide range of molecular dating methods available in the software.

We use both realistic simulations and empirical datasets of 40 Cyanobacteria and 62 Archaea to evaluate our method. We show that the combination of relative age constraints with fossil calibrations significantly improves the estimation of node ages.

## Introduction

Dated species trees (chronograms or timetrees, in which branch lengths are measured in units of geological time) are used in all areas of evolutionary biology. Their construction typically involves collecting molecular sequence data, which are then analyzed using probabilistic models. Commonly, a *relaxed molecular clock* approach is adopted. Such methods combine three components: a model of sequence evolution, a model of clock rate variation across the phylogeny, and calibrations of node ages based on the geological record. Inference is typically performed using Bayesian Markov chain Monte Carlo (MCMC) algorithms.

Inferring the age of speciations based on molecular data is challenging because it amounts to factoring divergence between sequences, estimated in units of substitutions per site, as a product of time (ages of splits) and rates of evolution. Additional information on ages and clock rates must be provided. Information on node ages is provided through *calibrated* nodes, *i.e*. nodes that can be associated to a date in the past, usually with some uncertainty. Node age calibrations are often derived from the ages of particular fossils or groups of fossils, but any information about dates in the past that can be associated with nodes (e.g., geochemical information such as the amount of oxygen in the atmosphere) can be used. By contrast, external data is rarely available to inform clock rates, especially over longer timescales where contemporary mutation rates, even if they are known, are not informative.

Consequently, inferences of the rate of evolution combine information contained in the analyzed sequence data and in the node age calibrations and are strongly dependent on the model of rate evolution along the phylogeny. When rates can be considered to be constant throughout the phylogeny, *i.e*. when the strict molecular clock hypothesis (Zuckerkandl and Pauling 1962) can be applied, only a single global rate needs to be estimated. For data sets that do not fit the strict molecular clock hypothesis, different rates need to be used to model sequence evolution in different parts of the tree. Several such relaxed clock models have been proposed (Thorne, Kishino, and Painter 1998; Drummond et al. 2006; Heath, Holder, and Huelsenbeck 2012; Lepage et al. 2007; Lartillot, Phillips, and Ronquist 2016) to account for rate variation across the phylogeny. Some assume that branch-specific rates are drawn independently of each other from a common distribution with global parameters (Drummond et al. 2006; Lepage et al. 2007; Heath, Holder, and Huelsenbeck 2012). Other models assume neighboring branches to have more similar rates than distant branches (Thorne, Kishino, and Painter 1998), and a model that can accommodate both situations has recently been proposed (Lartillot, Phillips, and Ronquist 2016). The sophistication, and typically much better fit (Pybus 2006) of relaxed clock models, however, comes at a price: inference is computationally more demanding than under the strict molecular clock. This is because relaxed clock models contain a large number of parameters, some of which are highly correlated.

Since the inference of the rate of evolution extracted by relaxed clock models contains uncertainty, dating a phylogeny relies heavily on the calibrations that are used to anchor the nodes in time (Pybus 2006; dos Reis, Donoghue, and Yang 2015). Unfortunately, fossils are rare and unevenly distributed both in the geological record and on the tree of life. Microbes, in particular, have left few fossils that can be unambiguously assigned to known species or clades. Therefore, entire clades cannot be reliably dated because they lack such information. For example, a recent dating analysis encompassing the three domains of life (Betts et al. 2018) used 11 fossil calibrations, 7 of which could be assigned to Eukaryotes, 3 to Bacteria, and none to Archaea. Clearly, incorporating new sources of information into dating analyses would be very useful, especially for dating the microbial tree of life.

Recently, it has been shown that gene transfers encode a novel and abundant source of information about the temporal coexistence of lineages throughout the history of life (Szöllosi et al. 2012; Davín et al. 2018; Wolfe and Fournier 2018; Magnabosco et al. 2018). From the perspective of divergence time estimation, gene transfers provide *node order constraints, i.e*., they specify that a given node in the phylogeny is necessarily older than another node, even though the older node is not an ancestor of the descendant node (Fig. 1a). Davín et al. (2018) showed that the dating information provided by these constraints was consistent with information provided by (calibrated) relaxed molecular clocks, which suggests that node calibrations could be combined with node order constraints to date species trees more accurately. The benefit of including transfer-based constraints may be particularly noticeable in microbial clades, where transfers can be frequent (Doolittle 1999; Abby et al. 2012; Szöllosi et al. 2012; Davín et al. 2018) and fossils are rare. However, constraints may also be derived from other events, such as the transfer of a parasite or symbiont between hosts, endosymbioses, or other obligatory relationships.

**Fig. 1.**
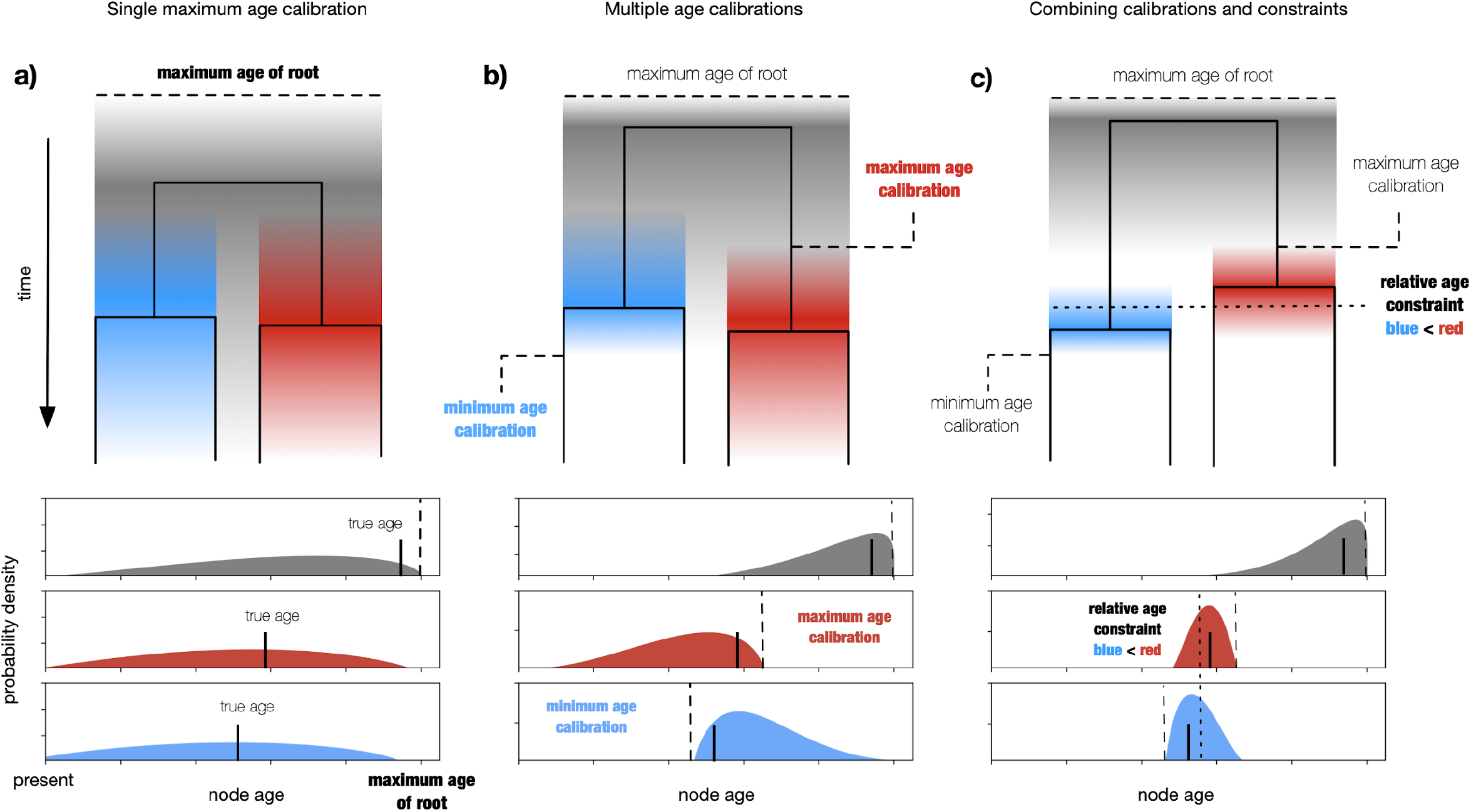
Relative age constraints inform molecular clock based dates. a) estimation of divergence time from sequences requires at least one maximum age calibration, typically provided as a maximum age of the root. As illustrated above with only a single maximum age calibration, the estimates will be highly uncertain. b) incorporating multiple minimum and maximum age calibrations, usually based on fossils from the geological record, can increase the resolution and accuracy of node ages, but well-resolved and accurate ages require large numbers of calibrations that are not always available. c) incorporating relative age constraints that specify that a given node in the phylogeny is necessarily older than another node, even though the older node is not an ancestor of the descendant node, can further improve the resolution and accuracy of molecular clock inferences.

The inclusion of relative age constraints into dating methods has so far involved ad-hoc approaches, involving several steps (Davín et al. 2018; Wolfe and Fournier 2018; Magnabosco et al. 2018). A statistically correct two-step approach was proposed by Magnabosco et al. (2018). First, an MCMC chain is run with calibrations but without relative age constraints. Then the posterior sample of timetrees is filtered to remove timetrees that violate relative age constraints. This approach works for a small number of constraints, but is difficult to scale to large numbers of constraints, where an increasing proportion of sampled timetrees will be rejected. Here, we present a method to combine node age constraints with node age calibrations within the standard (relaxed) molecular clock framework in a Bayesian setting. The resulting method is statistically sound and can handle a large number of constraints. We examine its performance on realistic simulations and evaluate its benefits on two empirical data sets.

## Materials and Methods

### Bayesian MCMC dating with calibrations and constraints

#### Informal description

Relaxed clock dating methods are often implemented in a Bayesian MCMC framework. Briefly, prior distributions are specified for (1) a diversification process (e.g., a birth-death prior), (2) the parameters of a model of sequence evolution (e.g., the HKY model, Hasegawa, Kishino,and Yano 1985), (3) calibration ages, and (4) the parameters of a model of rate heterogeneity along the tree. Such models may consider that neighboring branches have correlated rates of evolution (e.g., the autocorrelated lognormal model, Thorne, Kishino, and Painter 1998), or that each branch is associated to a rate drawn from a shared distribution (e.g. the uncorrelated gamma model Drummond et al. 2006). Calibrations specify prior distributions that account for the uncertainty associated with their age (dos Reis, Donoghue, and Yang 2015), and sometimes for the uncertainty associated with their position in the species tree (Heath, Huelsenbeck, and Stadler 2014). Our method introduces relative node age constraints as a new type of information that can be incorporated into this framework.

We chose to treat relative node order constraints as data without uncertainty, in the same way that topological constraints have been implemented in e.g. MrBayes (Ronquist and Huelsenbeck 2003). Note, our approach disregards uncertainty and differs from common node age calibrations.This decision provides us with a simple way to incorporate constraints in the model: during the MCMC, any tree that does not satisfy a constraint is given a prior probability of 0, and is thus rejected during the Metropolis-Hastings step. Therefore, only trees that satisfy all relative node age constraints have a non-zero posterior probability.

### Formal description

Let *A* be the sequence alignment, Ca be the set of fossil calibrations, and Co be the set of node order constraints. Further, let Ψ_*t*_ be the timetree, i.e. a tree with branch lengths in units of time (e.g. years), and Ψ_*s*_ be the tree with branches measured in expected number of substitutions per unit time, respectively. Finally, let *θ* be the set of all other parameters. In particular, *θ* contains the parameters of the sequence evolution model, the parameters of the relaxed molecular clock model, and the rates of the timetree diversification model. The sets (*A*, Co, Ca) and (Ψ_*s*_, Ψ_*t*_, *θ*) fully specify the data and the model, respectively. Then, the posterior distribution is

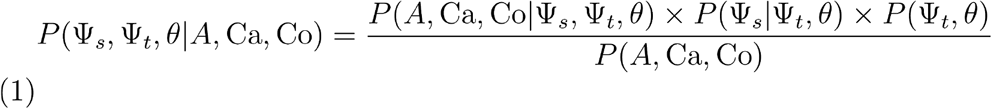

The likelihood consists of two terms, the first of which can be further separated into

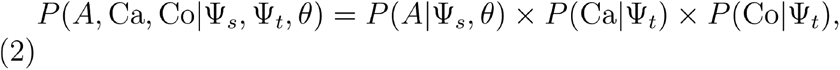

where *P*(*A*|Ψ_*s*_, *θ*) is the phylogenetic likelihood typically obtained with the pruning algorithm (Felsenstein 1981). The probability density *P*(Ca|Ψ_*t*_) assures the node age calibrations Ca are honored by Ψ_*t*_ using distributions with hard or soft boundaries (Yang and Rannala 2005). Node order constraints are accounted for by *P*(Co|Ψ_*t*_) = *δ*(Co, Ψ_*t*_), where *δ*(Co, Ψ_*t*_) is the indicator function that is one if the node order constraints Co are satisfied by Ψ_*t*_, and zero otherwise. The second term *P*(Ψ_*s*_|Ψ_*t*_, *θ*) of the likelihood in Equation (1) describes the relaxed molecular clock model, which includes the rate modifiers relating the branches in expected number of substitutions of Ψ_*s*_ to the branches in units of time of Ψ_*t*_. Here, we use the uncorrelated gamma relaxed molecular clock model, but many other models such as the lognormal relaxed molecular clock model are available (Lepage et al. 2007).

Finally, the prior *P*(Ψ_*t*_, *θ*) is usually separated into a product of a timetree prior *P*(Ψ_*t*_|*θ*) typically based on the birth-death process (Rannala and Yang 1996) and a prior *P*(*θ*) on the other parameters.

### Two-step inference of timetrees

Evaluation of the phylogenetic likelihood *P*(*A*|Ψ_*s*_, *θ*) in Equation (2) is the most expensive operation when calculating the posterior density. Further, the phylogenetic likelihood has to be recalculated at each iteration when performing a Bayesian MCMC analysis. Typically, the Markov chain has to be run for many iterations to obtain a good approximation of the posterior distribution. Consequently, inference is cumbersome, even when the topology of Ψ_*s*_ is fixed. To reduce the computational cost, we decided to approximate the phylogenetic likelihood using a two-step approach.

In the first step, the posterior distribution of branch lengths measured in expected number of substitutions is obtained for the fixed unrooted topology of Ψ_*s*_ using a standard MCMC analysis. The obtained posterior distribution is used to calculate the posterior mean *μ_i_* and posterior variance *υ_i_* of the branch length for each branch *i* ∈ *I* of the unrooted topology of Ψ_*s*_.

In the second step, the posterior means and variances are then used to approximate the phylogenetic likelihood using a composition of normal distributions

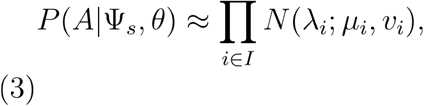

where λ_*i*_, which is sampled during this second MCMC analysis, is the branch length measured in expected number of substitutions of branch *i* of the unrooted topology of Ψ_*s*_. *N*(*x*; *μ*, *υ*) is the probability density of the normal distribution with mean *μ* and variance *υ* evaluated at *x*. Since the two branches leading to the root of Ψ_*t*_ correspond to a single branch on the unrooted topology of Ψ_*s*_, only their sum contributes to *P*(*A*|Ψ_*s*_, *θ*).

The two-step approach is similar to the approximation of the phylogenetic likelihood performed by MCMCTree (Reis and Yang 2011). MCMCTree uses a variable transformation together with a second order Taylor expansion of the likelihood surface, thereby also handling the covariance of branch lengths. The two-step approach reported here is fast for large data sets as well as complex models. In fact, state-of-the-art substitution models such as the CAT model, which is currently available only in PhyloBayes (Lartillot et al. 2013), could be used during the first step of the analysis.

### Implementation

We implemented this model and the two-step approach in RevBayes so that it can be combined with other available relaxed molecular clock models and models of sequence evolution and species diversification. Using the model in a RevScript implies calling two additional functions: one to read the constraints from a file, and another one to specify the timetree prior accounting for the constraints. Scripts are available at https://github.com/Boussau/DatingWithConsAndCal. We also provide a tutorial to guide RevBayes users: https://revbayes.github.io/tutorials/relative_time_constraints/

### Evaluation of the accuracy of the two-step approach

We compared our two-step, composite-likelihood approach to the one-step, full Bayesian MCMC approach in combination with two different models of rate evolution, White Noise (WN), and Uncorrelated Gamma (UGAM) (see Lepage et al. 2007 for a presentation of both). Analyses were performed in RevBayes (Höhna et al. 2016). We used an empirical sequence alignment and phylogeny of 36 mammalian Species from Reis et al. (2012), using all their calibrations and no relative constraint.

### Simulations to evaluate the usefulness of relative node age constraints

#### General framework

We generated an artificial timetree. We gathered calibration points by recording true node ages in this artificial timetree. We also gathered relative constraints by recording true relative orders between the nodes. Then we altered the branch lengths of the timetree to obtain branch lengths in expected number of substitutions (see Fig. S1 for a description of our simulation protocol). Based on this substitution tree, we simulated a DNA sequence alignment. Based on this sequence alignment, we used the two-step approach described above in RevBayes to infer timetrees. We then compared the reconstructed node ages to the true node ages from the artificial timetree to investigate the information provided by constraints.

#### Simulating an artificial timetree

To obtain a tree with realistic speciation times, we decided to simulate a tree that has the same speciation times as in the timetree of life from Betts et al. (2018). To do so, we gathered the speciation times from that timetree and produced an artificial tree by firstly randomly joining tips to produce speciation events, and secondly assigning the speciation times from the empirical timetree to these speciation events. We call the resulting tree a “shuffled tree” (Fig. 2). This shuffled tree has total depth from root to tips 45.12 units of time, as the timetree of life from Betts et al. (2018).

**Figure 2:**
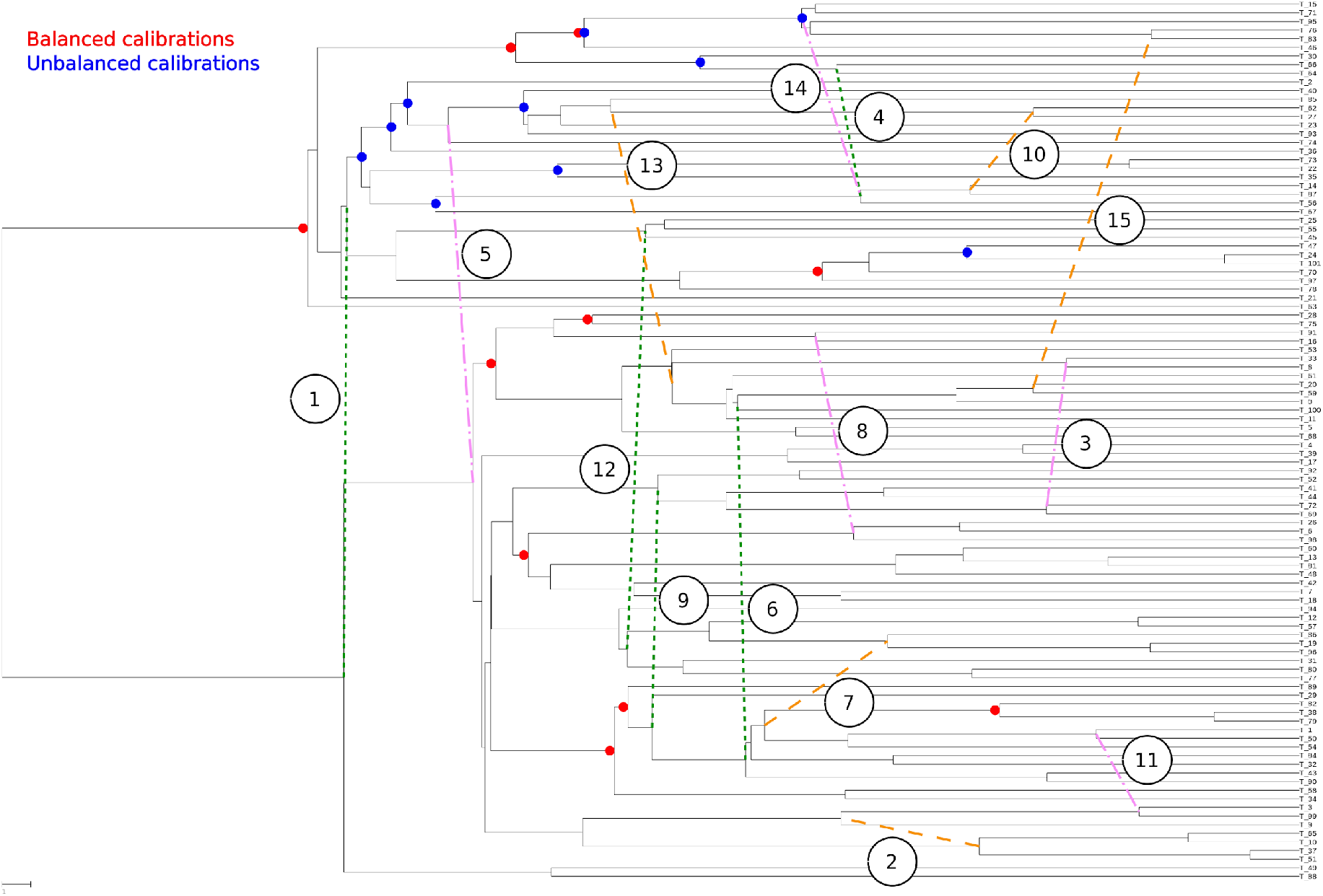
Shuffled tree, calibrated nodes and node order constraints. Calibrated nodes are shown with red dots when they are part of the set of 10 balanced calibrations, and with blue dots when they are part of the set of 10 unbalanced calibrations. Handpicked constraints have been numbered from 1 to 15, according to one order in which they were used (e.g. constraint 1 was used when only one constraint was included, constraints 1 to 5 when 5 constraints were included, and so on). Constraints have been colored according to their characteristics: green constraints are the 5 constraints between nodes with most similar ages (proximal), orange constraints are the 5 constraints between nodes with least similar ages (distal), and purple constraints are in between.

#### Building calibration times and node order constraints

We chose to use 10 internal node calibrations plus one calibration at the root node, as in Betts et al. (2018). We used two configurations: one *balanced* configuration where calibrations are placed on both sides of the root, and one *unbalanced* configuration where calibrations are found only on one side of the root (Fig. 2, red and blue dots respectively).

We selected 15 constraints by gathering true relative node orders from the shuffled tree. In choosing our sets of constraints we avoided redundant constraints, *i.e*. constraints that were already implied by previously included constraints. These constraints are shown Fig. 2. We performed one inference with 0 constraint, and one inference with all 15 constraints. In addition, we ran 10 independent experiments. In each experiment, we performed inference 14 times, varying the number of constraints from 1 to 14. The order with which constraints were introduced varied between experiments.

We built calibration times from the artificial tree by gathering the true speciation time, and associating it a prior distribution to convey uncertainty. The prior distribution we chose is uniform between [true age - (true age/5); true age + (true age/5)] and decays according to the tails of a normal distribution with standard deviation 2.5 beyond these boundaries (with 2.5% of the prior weight in each tail). 10 calibration points were chosen both in the balanced and unbalanced cases (Fig. 2). In addition, the tree root age was calibrated with a uniform distribution between [root age - (root age/5); root age + (root age/5)].

#### Simulations of deviations from the clock

The shuffled tree was rescaled to yield branch lengths that can be interpreted as numbers of expected substitutions (its length from root to tip was 0.451). Then it was traversed from root to tips, and rate changes were randomly applied to the branches according to two Poisson processes, one for small and frequent rate changes, and one for big and rare rate changes. The magnitudes of small and large rate changes were drawn from lognormal distributions with parameters (mean=0.0, variance=0.1) and (mean=0.0, variance=0.2), respectively, and their rates of occurrence were 33 and 1, respectively. After this process, branches smaller than 0.01 were set to 0.01. A Python code using the ete3 library (Huerta-Cepas, Serra, and Bork 2016) is available at https://github.com/Boussau/DatingWithConsAndCal/blob/master/Scripts/alterBranchLengths.py, along with the command lines used and a plot of the resulting tree with altered branch lengths (https://github.com/Boussau/DatingWithConsAndCal/blob/master/SimulatedTrees/proposedTree_rescaled_altered.dnd.pdf). The trees at the various steps of this simulation pipeline are also represented Fig. S1.

We compared the extent of the deviations from ultrametricity we had introduced in our simulated tree to empirical trees from the Hogenom database (Penel et al. 2009). Fig. S2 shows that our simulated tree harbours a realistic amount of non-ultrametricity.

#### Alignment simulation

The tree rescaled with deviations from the clock was used to simulate one alignment 1000 bases long according to a HKY model (Hasegawa, Kishino, and Yano 1985), with ACGT frequencies {0.18, 0.27, 0.33, 0.22} and with a transition/transversion ratio of 3. The Rev script is available at https://github.com/Boussau/DatingWithConsAndCal/Scripts/simu_HKY_No_Gamma.Rev.

#### Inference based on simulated data

Inference of timetrees based on the simulated alignment was performed in two steps as explained above. Both steps were performed in RevBayes (Höhna et al. 2016).

We inferred branch length distributions under a Jukes-Cantor model (Jukes and Cantor 1969) to make our test more realistic in that the reconstruction model is simpler than the process generating the data. The tree topology was fixed to the true unrooted topology. The script is available at https://github.com/Boussau/DatingWithConsAndCal/blob/master/Scripts/mcmc_JC.Rev.

The obtained posterior distributions of branch lengths were then summarized by their mean and variance per branch (script available at https://github.com/Boussau/DatingWithConsAndCal/blob/master/Scripts/DatingRevScripts/computeMeanAndVarBl.Rev). These means and variances were given as input to a script that computes a posterior distribution of timetrees according to a birth-death prior on the tree topology and node ages, an uncorrelated Gamma prior on the rate of sequence evolution through time (Drummond et al. 2006), and using the calibrations and constraints gathered in previous steps (see above), with the Metropolis Coupled Markov Chain Monte Carlo algorithm (Altekar et al. 2004). The corresponding script is available at https://github.com/Boussau/DatingWithConsAndCal/blob/master/Scripts/DatingRevScripts/mainScript.Rev.

#### Empirical data analyses

We used alignments, tree topologies and sets of constraints from Archaea and Cyanobacteria that had been previously analyzed in Davín et al. (2018). In both cases, the constraints had been derived from transfers identified in the reconciliations of thousands of gene families with the species tree, and filtered to keep the largest consistent set of supported constraints. We used 431 constraints for Archaea, and 144 for Cyanobacteria. Alignments, trees and sets of constraints are available at https://doi.org/10.5061/dryad.s4mw6m958. We used the CAT-GTR model in Phylobayes (Lartillot et al. 2013) to generate branch length tree distributions with a fixed topology, and our two-step approach in RevBayes (Höhna et al. 2016) to compute posterior distributions of timetrees, under the UGAM model of rate evolution (Drummond et al.2006).

## Results

### Two-step inference provides an efficient and flexible method to estimate time trees

We compared posterior distributions of node ages obtained using the classical full Bayesian MCMC approach to those obtained using our two-step approximation on a dataset of 36 mammalian species (dos Reis et al. 2012). As shown in Supplementary Figs. S3-6, the two posterior distributions of node ages are practically indistinguishable. Further, the impact of the approximation is negligible in comparison to the choice of the model of rate evolution. We used the uncorrelated Gamma (UGAM) or the White Noise (WN) models, both uncorrelated, and found that using one or the other results in more differences in the estimated node ages than using our two-step inference compared to the full Bayesian MCMC.

### Simulations

#### Constraints improve dating accuracy

We used two statistics to evaluate the accuracy of node age estimates. Firstly, we computed the normalized root mean square deviation (RMSD) between the true node ages used in the simulation and the node ages estimated in the Maximum A Posteriori tree (Fig. 2a), and normalized it by the true node ages. This provides measures of the error as a percentage of the true node ages. Secondly, we computed the coverage probability, *i.e*. how frequently the 95% High Posterior Density (HPD) intervals on node ages contained the true node ages (Fig. 2b).

As the number of constraints increases, Fig. 3a shows that the error in node ages decreases and Fig. 3b shows that the 95% HPD intervals include the true node ages more often. When 0 or only 1 constraint is used, the true node age is contained in only ~55% of the 95% HPD intervals, suggesting that the mismatch between the model used for simulation and the model used for inference has a noticeable impact. Poor mixing could also explain these results, but it is unlikely to occur in our experiment for two reasons. First, the Expected Sample Sizes for the node ages are typically above 300. Second, if the same moves are used in the MCMC, but the simulation model is changed to fit the inference model, about 95% of the true node ages end up in 95% HPD intervals, as expected for well-calibrated Bayesian methods and well-mixing MCMC chains (see Supp. Fig. S8 and associated section).

**Figure 3:**
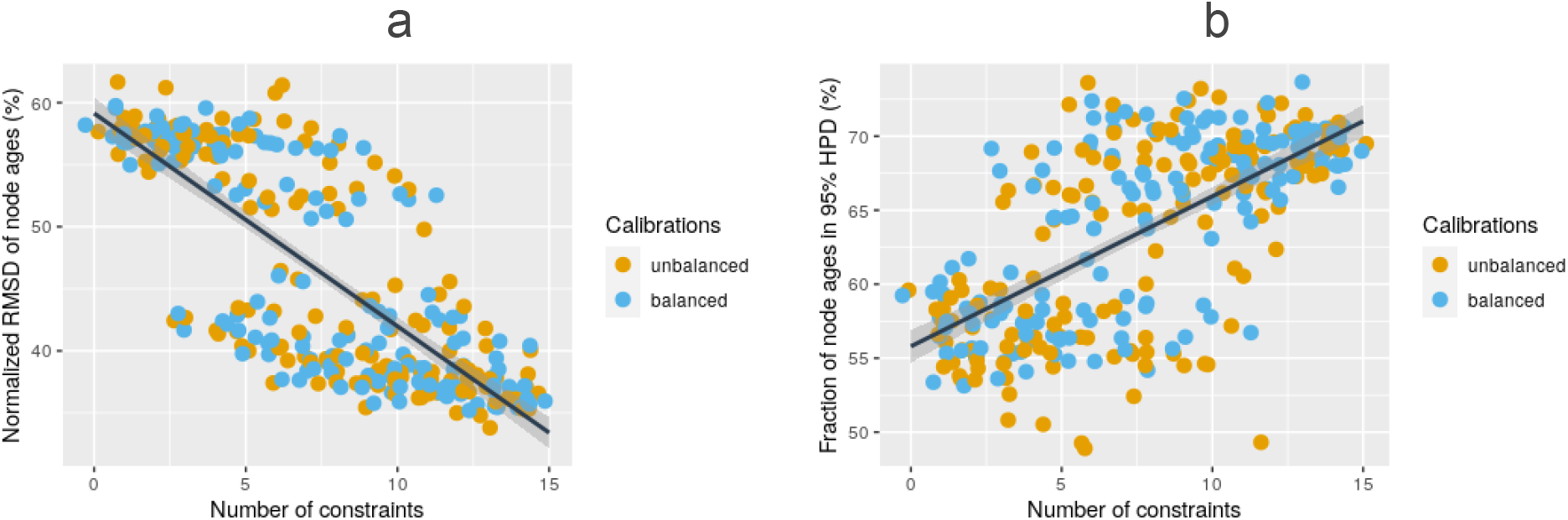
Increasing the number of constraints improves node age estimation. a) Average normalized RMSD over all internal node ages is shown in orange for 10 balanced calibrations and blue for 10 unbalanced calibrations. This is a measure of the error as a percentage of the true node ages. b) The percentage of nodes with true age in 95% High Posterior Density (HPD) interval is shown (colors as in a). Regression lines with confidence intervals in grey have been superimposed.

Results improve with more constraints. The variation in normalized RMSD can be explained by a linear model (M1) including an intercept and the number of constraints with an adjusted R-squared of ~0.63. However, it appears that points in Fig. 3a can be grouped in at least two clusters: those with normalized RMSD above ~48%, and those below. This suggests that some constraints have a bigger effect than other constraints. In particular, constraint 5 (see Fig. 2) is absent from all runs with normalized RMSD above 48%, suggesting that it is highly informative (more on the informativeness of constraints below).

The results obtained with the balanced set of calibrations are similar to the results obtained with the unbalanced set of calibrations: adding a variable indicating whether the balanced or unbalanced sets were used to model M1 does not improve the adjusted R-squared.

#### Constraints reduce credibility intervals

The additional information provided by constraints results in smaller credibility intervals, as shown in Fig. 4. The improvement in coverage probability observed in Fig. 3b therefore occurs despite smaller credibility intervals.

**Figure 4:**
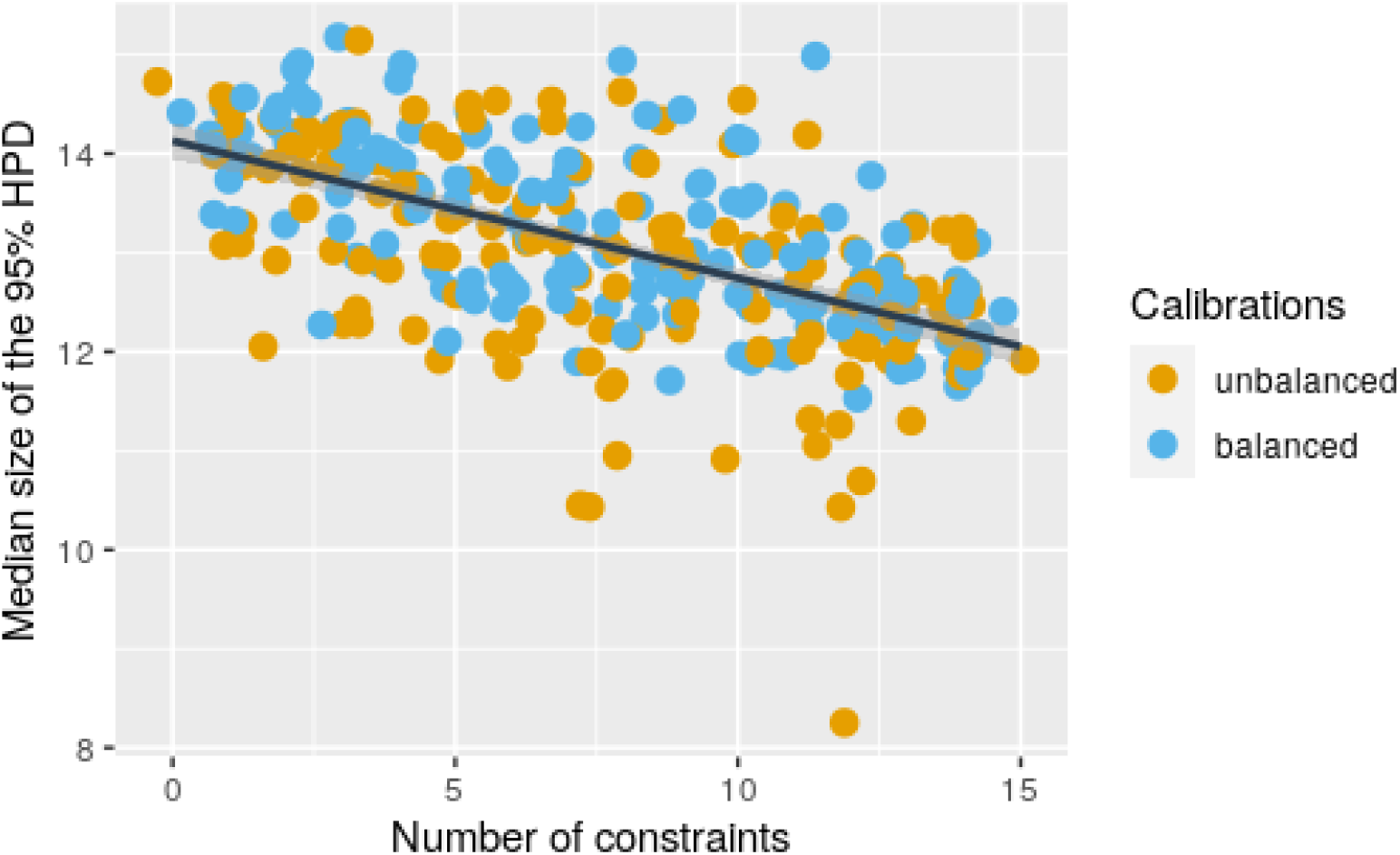
The 95% HPD intervals on node ages become smaller as the number of constraints increases. The sizes are given in units of time; for reference, the total depth for the true tree is 45.12 units of time. Colors as in Fig. 2. A regression line with confidence intervals in grey has been superimposed.

#### Investigating the informativeness of constraints

To measure the informativeness of constraints, we developed a linear model predicting the normalized RMSD based on whether or not each of the 15 constraints were used. This linear model improves upon M1 with an adjusted R-squared of 0.91. Its coefficients provide a measure of the informativeness of each constraint (Fig.5).

**Figure 5:**
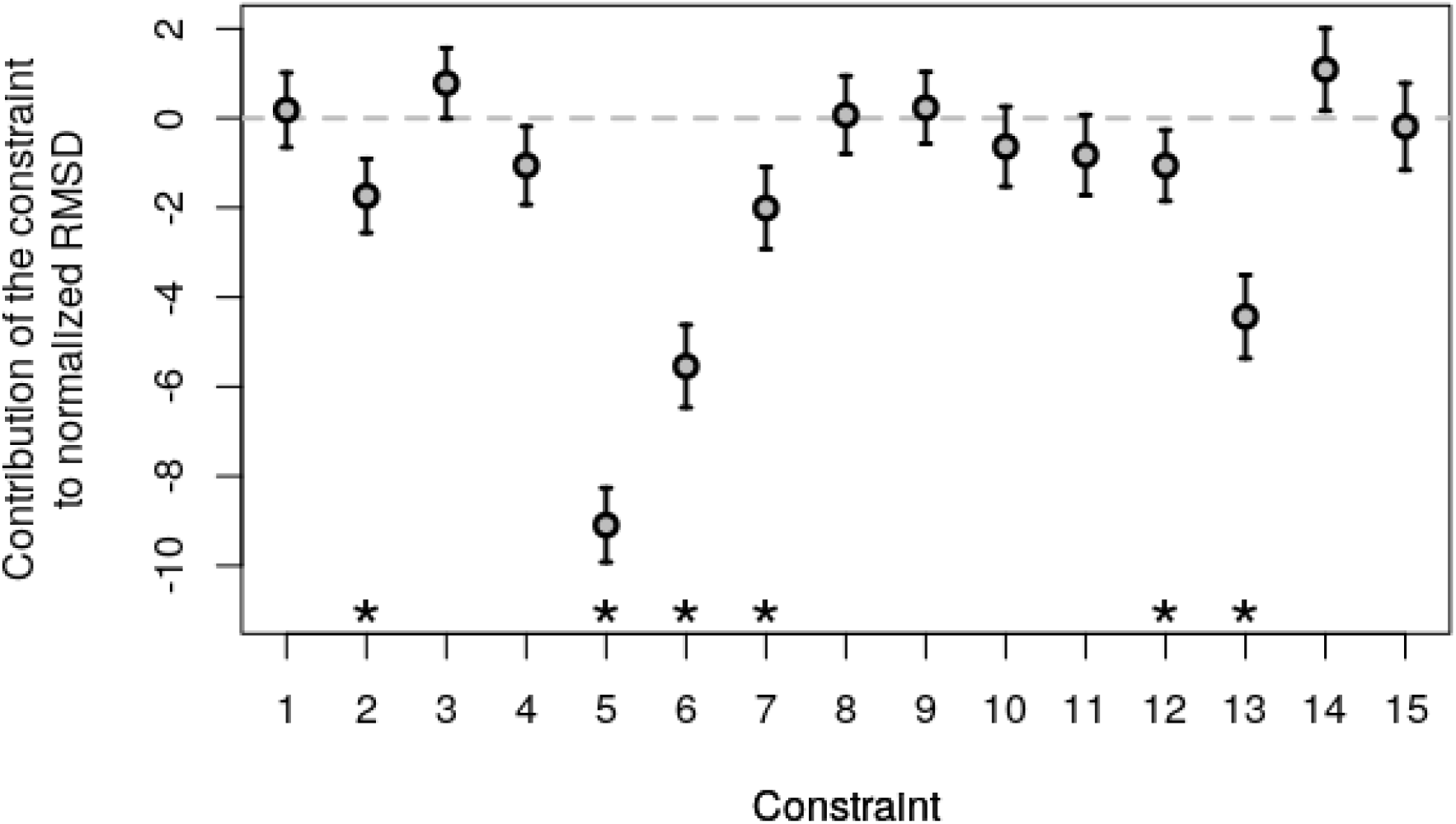
Contribution of individual constraints to dating error. Each constraint reduces up to 9.1 normalized RMSD percentage points. Error bars correspond to twice the standard error. Stars indicate coefficients of the linear model that are significantly different from 0 at the 1% level. Computations were run with 10 balanced or unbalanced calibrations.

Some constraints are much more informative than other constraints. Constraint 5 is the most informative one, as it reduces the normalized RMSD by 9.1 percentage points, followed by constraint 6, which reduces RMSD by 5.5 points, and constraint 13 which reduces RMSD by 4.4 points. All provide a significant reduction in normalized RMSD according to our linear model at the 1% level, along with constraints 2, 7 and 12. Constraints 1, 3, 4, 8, 9, 10, 11, 14 and 15 do not bring much information as they do not significantly affect the normalized RMSD at the 1% level. Constraints 3 and 14 appear to increase the normalized RMSD if the significance threshold is increased to 5%.

To understand what explains the difference in informativeness among our constraints, we computed statistics associated with each of them. We provide a more detailed discussion of what could make a constraint informative in the supplementary material, but here we investigated 8 different statistics computed on the true timetree. Firstly, three statistics computed between the two constrained nodes: the difference in true node ages, the nodal distance and the sum of branch lengths. We also noted whether the constraint spanned the root node, computed the number of leaves in the older and younger subtrees involved in the constraint, and the number of nodes ancestral to the nodes involved in the constraint. We regressed the contributions of each constraint to the normalized RMSD (Fig. 5) against these 8 statistics. We obtained an adjusted R-squared of ~0.67. The number of leaves in the younger subtree was the only significant explanatory variable at the 5% threshold, and the sum of branch lengths between the two constrained nodes came second (6.7%).

A constraint such that the younger node is the ancestor of a big subtree brings a lot of information because it provides an upper time constraint to all the nodes in the subtree. This is particularly useful in our context where all calibrations are lower time calibrations.

### Analyses of empirical data

Davín et al. (2018) showed that gene transfers contain dating information that is consistent with relaxed molecular clock models. We used a phylogeny of cyanobacterial genomes presented in Davin et al. and a phylogeny of archaeal genomes from Williams et al. (2017) to investigate the individual and cumulative impacts of fossil calibrations and relative constraints on the inference of time trees.

#### Relative constraints agree with fossil calibration on the age of akinete-forming multicellular Cyanobacteria

Davín et al. (2018) analyzed a set of 40 cyanobacteria spanning most of their species diversity. Cyanobacteria likely originated more than 2 billion years ago, but a review of the literature suggests that there is only a single reliable fossil calibration that we can place on the species tree: a minimum bound for akinete-forming multicellular Cyanobacteria from Tomitani et al. (2006). These authors reported a series of fossils that they assign to filamentous Cyanobacteria producing both specialized cells for nitrogen fixation (heterocysts) and resting cells able to endure environmental stress (akinetes).

We investigated whether relative node order constraints could recover the effect of the available fossil calibration by comparing several dating protocols: fossil calibration with no relative age constraints (Fig. 6a), no fossil calibration and no relative age constraints (Fig. 6b), relative age constraints with no fossil calibration, (Fig. 6c), and both calibrations and constraints (Fig. 6d). Fossil calibration corresponded to a minimum age for fossil akinetes at 1.956 GYa (dashed red line Arrow on Fig. 6a and d). Reflecting our uncertainty regarding the age of the root, we tried two alternatives for the maximum root age (*i.e*. age of crown cyanobacteria), 2.45 Gy and 2.7 Gy, corresponding to the “Great Oxygenation Event” and the “whiff of Oxygen” (Holland 2006) respectively. In Fig 6. we show results obtained with the 2.45 Gy root calibration, while Fig. 7a presents the age of key nodes for both choices of root maximum age.

**Figure 6.**
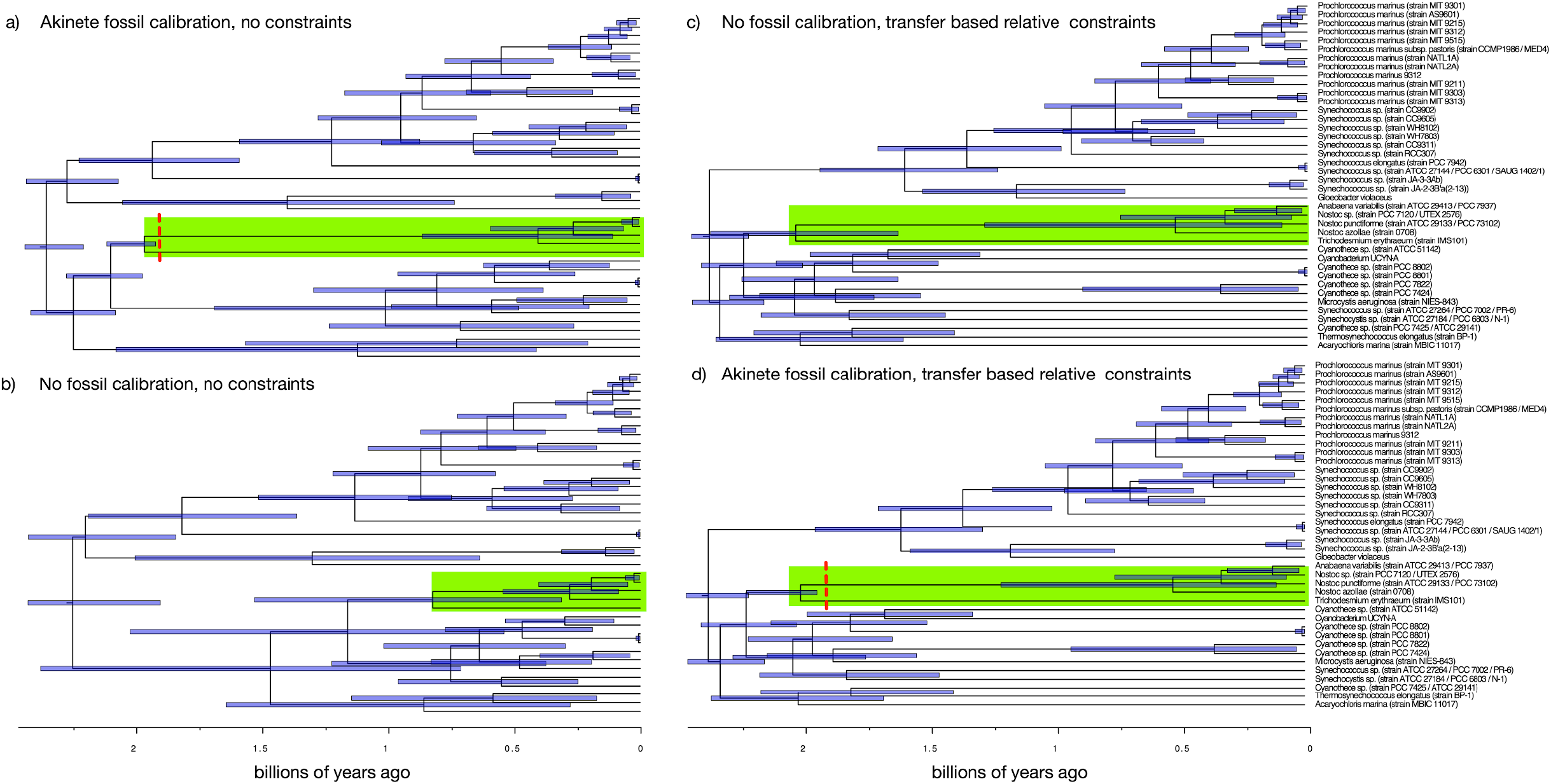
Relative age constraints agree with the akinete fossil calibration that akinete-forming multicellular Cyanobacteria are likely older than suggested by sequence data alone. We compared four dating protocols for the 40 cyanobacteria from Davin et al. (2018): a) fossil calibration (dashed red line) with no relative age constraints, b) no fossil calibration and no relative age constraints, c) 144 relative age constraints, with no fossil calibration and d) simultaneous fossil calibration and constraints (Fig. 5d). All four chronograms were inferred with a root maximum age of 2.45 Gya with an uncorrelated gamma rate prior, and a birth-death prior on divergence times. Clade highlighted in green corresponds to akinete-forming multicellular cyanobacteria.

**Figure 7:**
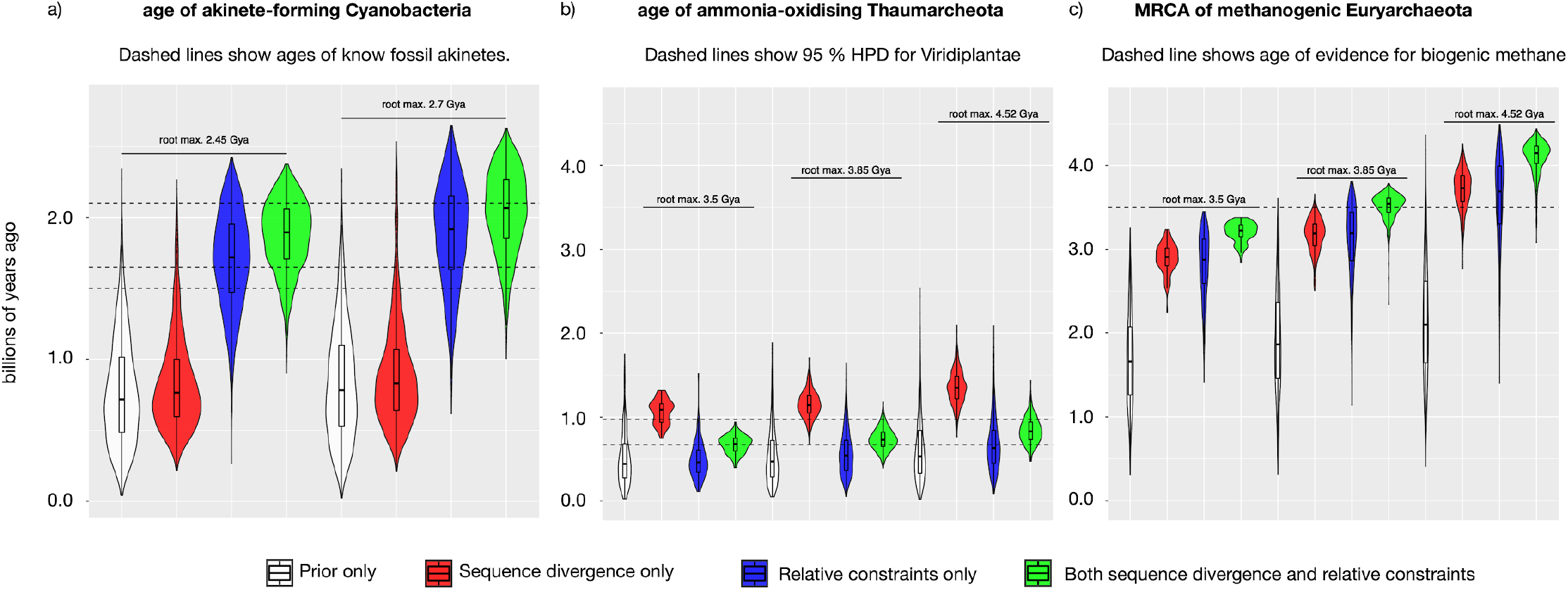
Distributions of key node ages according to different sources of dating information. We show the age of a) akinete-forming Cyanobacteria, b) Thaumarchaeota and c) the most recent common ancestor of methanogenic Archaea. Distributions in white are based on solely the maximum root age and the rate and divergence time priors, distributions in red are informed by sequence divergence, distributions in blue include relative age constraints, but not sequence divergence, while distributions in green rely on both. Dashed lines indicate, respectively, a) age of fossils of putative akinete forming multicellular cyanobacteria, b) age of Viridiplantae and c) age of evidence for biogenic methane. For the corresponding time trees with constraints see Supplementary Figure 9.

Comparison between Figs. 6a and 6b shows that including the minimum calibration increases the age of the clade containing akinete-forming multicellular Cyanobacteria (green clade) by about 1 Gy. Interestingly, the inclusion of constraints compensates for the absence of a minimum calibration (Fig. 6c) and places the age of clade of akinete forming multicellular Cyanobacteria close to its age when a fossil-based minimum age calibration is used (Fig.6a) calibrations are used. The information provided by constraints thus agrees with the fossil age for multicellular Cyanobacteria. As a result, the combination of both calibrations and constraints produces a chronogram with smaller credibility intervals (Fig. 6d).

To further characterize the effect of constraints on the age of akinete forming multicellular Cyanobacteria, we plotted the distributions of its age based on different sources of dating information. In Figure 7a we show the age of akinete forming multicellular Cyanobacteria (green clade in Fig 6) estimated based on i) only the rate and divergence time priors, ii) priors and sequence divergence only, iii) priors and relative age constraints only, and iv) priors and both sequence divergence and relative age constraints. Comparison of the age distributions shows that relative age constraints convey information that complements sequence divergence and is coherent with the fossil record on the age of akinete-forming Cyanobacteria.

#### Relative constraints refine the time tree of Archaea

We next investigated divergence times of the Archaea, one of the primary domains of life (Woese, Kandler, and Wheelis 1990). We used the data from Williams et al. (2017) containing 62 species. Most analyses place the root of the entire tree of life between Archaea and Bacteria (Woese, Kandler, and Wheelis 1990; Iwabe et al. 1989; Gogarten et al. 1989; Gouy, Baurain,and Philippe 2015), suggesting that the Archaea are likely an ancient group. However, there are no unambiguous fossil Archaea and so the history of the group in geological time is poorly constrained. Methanogenesis is a hallmark metabolism of some members of the Euryarchaeota, and so the discovery of biogenic methane in 3.46Gya rocks (Ueno et al. 2006) might indicate that Euryarchaeota already existed at that time. However, the genes required for methanogenesis have also been identified in genomes of other archaeal groups including Korarchaeota (McKay et al. 2019) and Verstraetearchaeota (Vanwonterghem et al. 2016), and it is difficult to exclude the possibility that methanogenesis maps to the root of the Archaea (Berghuis et al. 2019). Thus, ancient methane might have been produced by Euryarcheota, another extant archaeal group, a stem archaeon or even by Cyanobacteria (Bižić et al. 2020).

In the absence of strong geochemical constraints, can relative constraints help to refine the time tree of Archaea? We investigated two nodes on the archaeal tree from Williams et al. (2017): the common ancestor of ammonia-oxidising (AOA) Thaumarchaeota and the common ancestor of methanogenic Euryarchaeota (that is, the common ancestor of all Euryarchaeota except for the Thermococcus/Pyrococcus clade). While we lack absolute constraints for these lineages, dating hypotheses have been proposed on the basis of individually identified and curated gene transfers to, or from, other lineages for which fossil information does exist. These include the transfer of a DnaJ-Fer fusion gene from Viridiplantae (land plants and green algae) into the common ancestor of AOA Thaumarchaeota (Petitjean et al. 2012), and a transfer of three SMC complex genes from within one clade of Euryarchaeota (Methanotecta, including the class 2 methanogens) to the root of Cyanobacteria (Wolfe and Fournier 2018). Note that, in the following analyses, we did not use the two transfers listed above. Instead, we used 431 relative constraints derived from inferred within-Archaea gene transfers; therefore, these constraints are independent of the transfers used to propose the hypotheses we test.

As the age of the root of Archaea is uncertain, we explored the impact on our inferences of three different choices: a relatively young estimate of 3.5Gya from the analysis of Wolfe and Fournier (2018); the end of the late heavy bombardment at 3.85Gya (Boussau and Gouy 2012); and the age of the solar system at 4.52Gya (Barboni et al. 2017).

We found that, despite the uncertainty in the age of the root, the estimated age of AOA Thaumarchaeota informed by relative age constraints is consistent with the hypothesis that AOA are younger than stem Viridiplantae (Petitjean et al. 2012), with a recent estimate for the age of Viridiplantae between 972.4-669.9 Mya (Morris et al. 2018); (Figure 7b). As in the case of Cyanobacteria, information from relative constraints had a substantial impact on the analysis; sequence data alone (in combination with the root age prior) suggest a somewhat older age of AOA Thaumarchaeota, consistent with recent molecular clock analyses (Ren et al. 2019).

In the case of methanogenic Euryarchaeota, inference both with and without relative constraints was strongly influenced by the choice of root prior (Figure 7c), and so the results do not clearly distinguish between hypotheses about the age of archaeal methanogenesis or the potential source of ancient biogenic methane. With those caveats in mind, the information from relative constraints supported moderately older age distributions than inference from sequence data alone across all root priors. The results are consistent with an early origin of methanogenic Euryarchaeota within the archaeal domain (Wolfe and Fournier 2018) and, for the moderate (3.85Gya) and older (4.52Gya) priors, indicate that these archaea are a potential source of biogenic methane at 3.46Gya (Ueno et al. 2006).

## Discussion

### Constraints are a new and reliable source of information for dating phylogenies

Davín et al. (2018) showed that gene transfers contained reliable information about node ages. They also used this information in an *ad hoc* two-step process to provide approximate age estimates for a few nodes in 3 clades. Here we built upon these results to develop a fully Bayesian method that accounts for both relative node order constraints and absolute time calibrations within the MCMC algorithm by extending the standard relaxed clock approach. We also introduced a fast and accurate two-step method for incorporating branch length distributions inferred under complex substitution models into relaxed molecular clock analyses.

To test our method, we performed sequence simulations and analyzed three empirical data sets. We simulated sequences according to a model that differs from the inference model so as to emulate the typical situation with empirical data, where the process that generated the data differs from our inference models. As expected under these conditions, node age coverage probabilities, *i.e*., the percentage of true node ages that fall within inferred 95% credibility intervals, are much lower than 95%. We used a realistic phylogeny for simulating sequences by drawing node ages from a previously published dated tree of life (Betts et al. 2018) but by rearranging the tree topology. We then investigated the effect of sampling node age and node relative order constraints on dating accuracy. A single tree topology and a single simulated alignment were used overall, which might adversely affect the generality of our results. However, this tree topology is large (102 tips) and realistic, and the results on empirical data suggest that our method is useful across the tree of life. Further, using a single alignment allowed us to estimate branch length distributions only once and then use our fast two-step inference to reduce our computational footprint.

The simulations show that relative node order constraints improve the accuracy of node ages and coverage probabilities. We further found that some constraints were more informative than others. In particular, constraints in which younger nodes were ancestral to lots of nodes tended to be more informative than other constraints. This is because such a constraint provides an upper time limit to all the nodes in the younger subtree, which is complementary to the calibrations that provide lower time limits in our test. In empirical data analyses, lower time calibrations are more frequent than upper time calibrations, which suggests that there too the most informative constraints are likely to involve younger nodes ancestral to a big subtree.

Results obtained on empirical data sets show that relative node order constraints extracted from dozens of gene transfers contain information that can compensate for the lack of fossil calibrations. This shows promise for dating phylogenies for which fossils are scant, *i.e*., the great majority of the tree of life.

One limitation of the method presented here is that relative constraints are treated as though they are known with certainty. Only trees that satisfy all of the input constraints will have non-zero probability, and so incorrect input constraints will result in incorrect age estimates. We, therefore, suggest that only the most reliable constraints should be used when dating a species tree using transfers. One practical approach, which we have used in our empirical analyses of genomic data, is to use only those constraints that are highly supported (Davín et al. 2018). A clear direction for future work will be to treat relative constraints probabilistically, perhaps as a function of the number and quality of inferred gene transfers that support them, or with a probability *p* that constraints are matched, which would be estimated in the course of the MCMC.

Dating phylogenies is a challenging statistical problem where data is limiting since only fossils and rates of molecular evolution provide information. Here we have developed a new method to exploit the information contained in gene transfers, which are particularly numerous in clades where fossil information is lacking. Gene transfers define relative node order constraints. We have shown in simulations that using node order constraints improves node age estimates and reduces credibility intervals. We have also used our method on two empirical data sets to show that node order constraints can compensate for the absence of a fossil calibration: ages obtained without a fossil calibration but with constraints match those obtained with the fossil calibration, and incorporating both sources of time information further refines the inferred divergence times. Looking forward we envision that our method will be useful to date parts of the tree of life where node ages have so far remained very uncertain.

## Supplementary material

Supplementary Material is available at BioRxiv: https://www.biorxiv.org/content/10.1101/2020.10.17.343889v8.supplementary-material

## Data availability

Scripts and data used to run the simulation analyses are available at https://github.com/Boussau/DatingWithConsAndCal

Data for the empirical data analysis has been deposited at: https://doi.org/10.5061/dryad.s4mw6m958

A tutorial is available at: https://revbayes.github.io/tutorials/relative_time_constraints/ to use both our two-step approach and for dating with relative node age constraints.

## Author contributions

GJS, VD and BB initiated the project. BB, GJS and SH implemented the model in RevBayes. GJS ran the empirical analyses, and analyzed them with TAW. BB ran the simulations. DS, GJS and BB wrote the tutorial. BB, GJS, TAW and VD wrote the manuscript. All authors read and commented on the manuscript.

## Acknowledgements

Version 8 of this preprint has been peer-reviewed and recommended by Peer Community In Evolutionary Biology (https://doi.org/10.24072/pci.evolbiol.100127). We thank the reviewers and the editor for their comments on earlier versions of the manuscript. DS and GJSz received funding from the European Research Council under the European Union’s Horizon 2020 research and innovation program under Grant Agreement 714774, GJS received funding Grant GINOP-2.3.2.-15-2016-00057. TAW is supported by a Royal Society University Research Fellowship and NERC grant NE/P00251X/1. BB and TAW acknowledge support from a “Projet de Recherche Collaborative” co-funded by the CNRS and the Royal Society. We thank Eric Tannier for fruitful discussions.

## Conflict of interest disclosure

The authors of this preprint declare that they have no financial conflict of interest with the content of this article. GJZ, TAW and VD are members of the PCI Evol Biol recommenders.

## Notes

https://doi.org/10.5061/dryad.s4mw6m958

https://github.com/Boussau/DatingWithConsAndCal

https://revbayes.github.io/tutorials/relative_time_constraints/

### Competing Interest Statement

The authors have declared no competing interest.

### Summary of Updates

Version 8 of this preprint has been peer-reviewed and recommended by Peer Community In Evolutionary Biology (https://doi.org/10.24072/pci.evolbiol.100127); this version improves on version 7 with a correction of the link to the supplementary material that was not correct, and a change of format to ensure that text is not transformed into a jpeg next to the title.

