## Supplementary material for "Relative time constraints improve molecular dating"

Gergely J. Szöllősi: MTA-ELTE "Lendület"' Evolutionary Genomics Research Group, Pázmány P. stny. 1A, H-1117 Budapest, Hungary; Department of Biological Physics, Eötvös University,Pázmány P. stny. 1A, H-1117 Budapest, Hungary; Institute of Evolution, Centre for Ecological Research, Konkoly-Thege M. út 29-33. H-1121 Budapest, Hungary

Sebastian Höhna: GeoBio-Center LMU, Ludwig-Maximilians-Universität München, Richard-Wagner Straße 10, 80333 Munich, Germany; Department of Earth and Environmental Sciences, Paleontology & Geobiology, Ludwig-Maximilians-Universität München, Richard-Wagner Straße 10, 80333 Munich, Germany. *Email:*

### Table of contents

**Table of contents** 1

**Simulation protocol** 2

**Variance in root-to-tip length of the simulated substitution tree** 3

**Two-step inference of timetrees provides results similar to the usual one-step dating method** 4

**On the informativeness of constraints** 8

**Verification that the MCMC dating algorithm is well calibrated.** **9**

**Timetrees for the Archaea for different maximum root ages** **10**

**Bibliography** **12**

### Simulation protocol

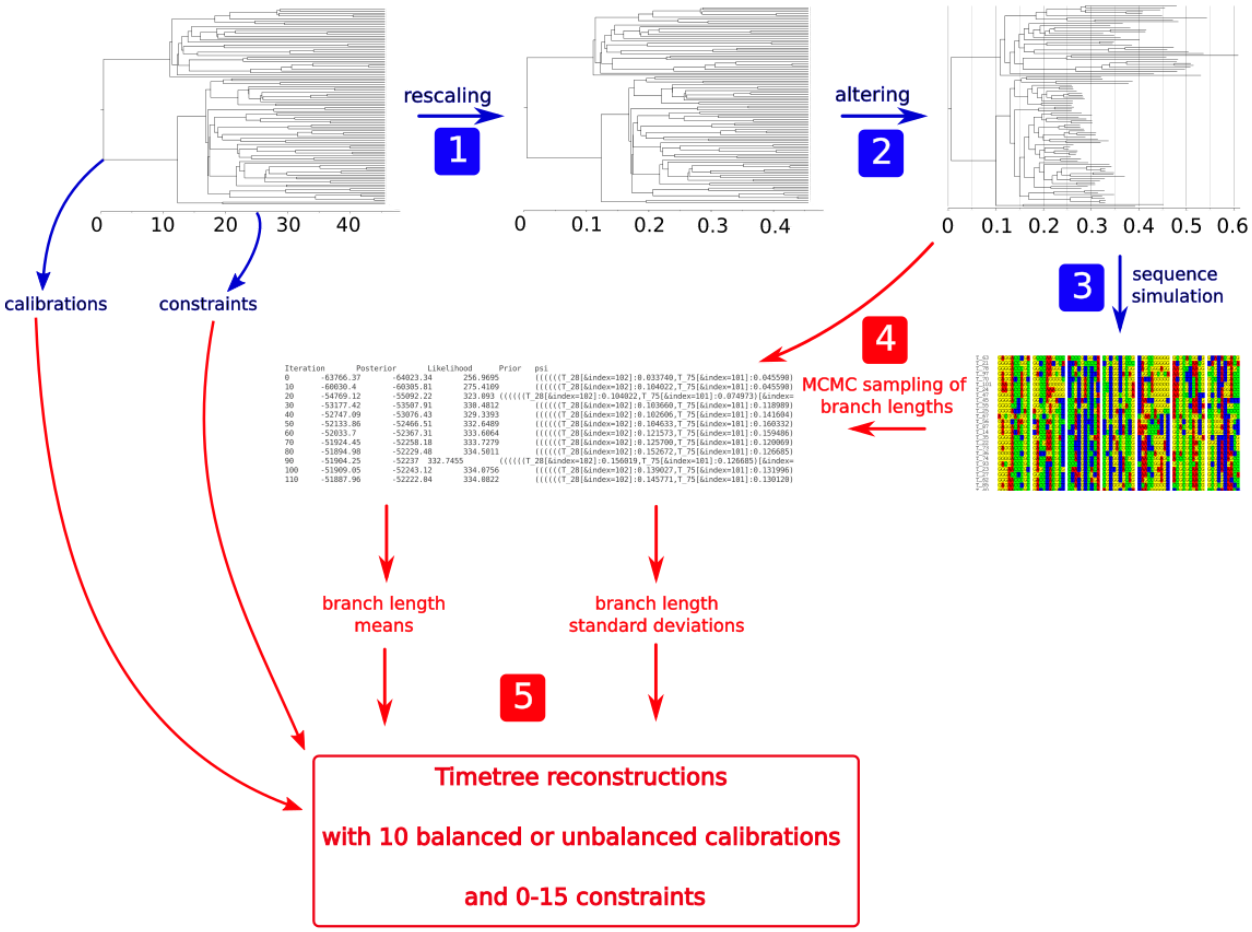


**Figure S1. Simulation and inference from simulated data**. The first three steps simulate an alignment, and the two last steps reconstruct timetrees in a variety of parameter settings. Starting top left from a time tree in units of time, several steps are applied to alter the branch lengths. 1, a rescaling is applied to change the branch lengths into expected numbers of substitutions. 2, branch lengths are randomly lengthened or shortened to simulate rate variation across lineages, resulting in what we call here a “substitution tree”. 3, an alignment is simulated along the obtained phylogeny according to a HKY model of sequence evolution with rate heterogeneity. 4, the alignment and the substitution tree are used as input into RevBayes, which generates a posterior sample of trees keeping the topology fixed. The branch length means and standard deviations are extracted from the sample. 5, several timetree reconstructions are performed, varying the number of constraints, their order, and using 10 balanced or unbalanced calibrations. The three trees that are shown are the trees that were used in our simulation.

### Variance in root-to-tip length of the simulated substitution tree

We compared the variance in root-to-tip length in our simulated substitution tree that was used to simulate sequences (after step 2, Fig. S1) to the same statistic computed on empirical substitution trees. We collected trees with branch lengths in events of substitutions from the Hogenom database v7 (Penel et al. 2009). We selected 121,182 trees that had been built from protein sequences of 666 genomes spanning the whole tree of life using IQTree (LG+PMSF+G model), and then had been midpoint-rooted. We further selected 1158 trees that had between 90 and 110 sequences from those trees.

For both the simulated substitution tree and the empirical trees, we computed root to tip distances. For each tree we normalized the distances by dividing them by the mean distance, and then computed the variance of these normalized distances. We plot the results Fig. S2.

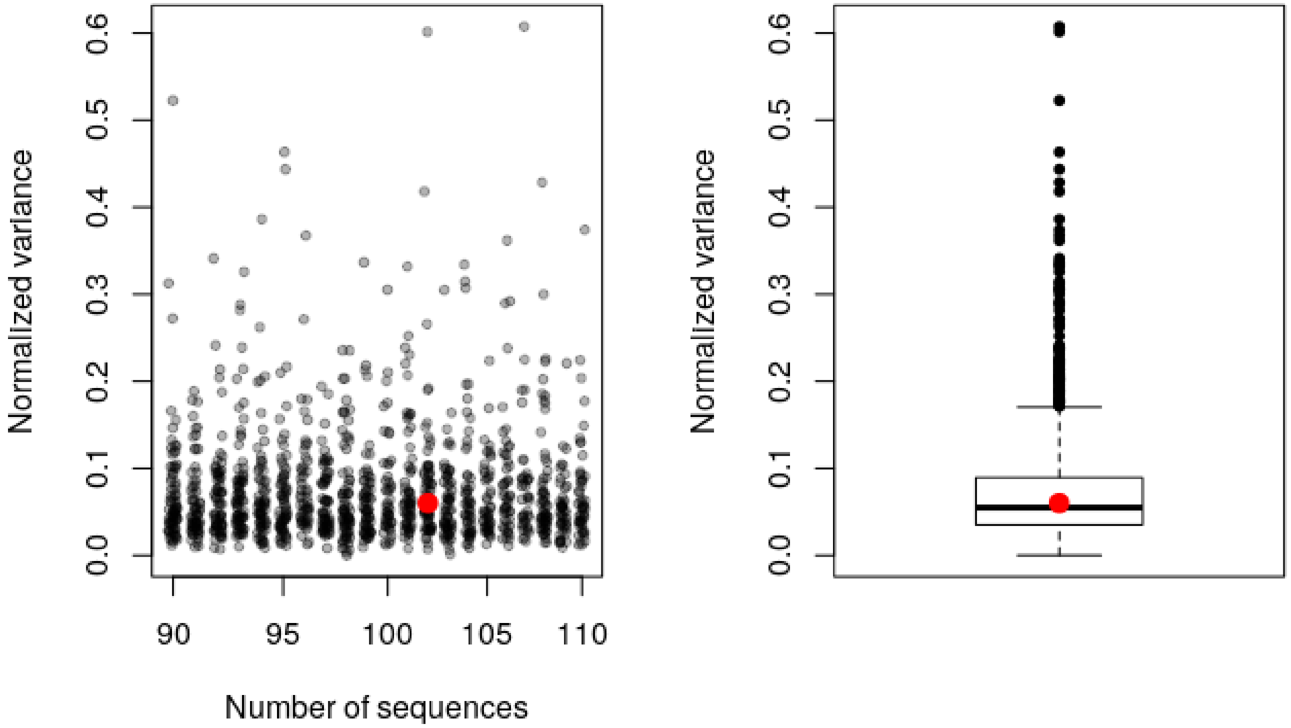

**Figure S2. Comparison of normalized variance in root-to-tip length between the simulated tree and empirical trees**. Left: Scatterplot of normalized root-to-tip length variances for all empirical trees between 90 and 110 sequences. A bigger red dot marks the value obtained for the simulated tree. Right: Boxplot representation of the same data.

The variance in root-to-tip length of our simulated tree is similar to that of empirical trees in the Hogenom database: 55% of our selection of empirical trees are more clockwise than our simulated tree.

### Two-step inference of timetrees provides results similar to the usual one-step dating method

We compared the effect of using our two-step, composite-likelihood approach instead of the one-step, full Bayesian MCMC approach, to that of using two different models of rate evolution, White Noise (WN), and Uncorrelated Gamma (UGAM) (see (Lepage et al. 2007) for a presentation of both). We used an empirical sequence alignment of 36 mammalian Species and a fixed tree topology from dos Reis et al. (dos Reis et al. 2012). Computations were run in triplicates. Results show that the effect of the model of rate evolution is much more important than that of using the two-step approach (Supp Fig. S3-S6).

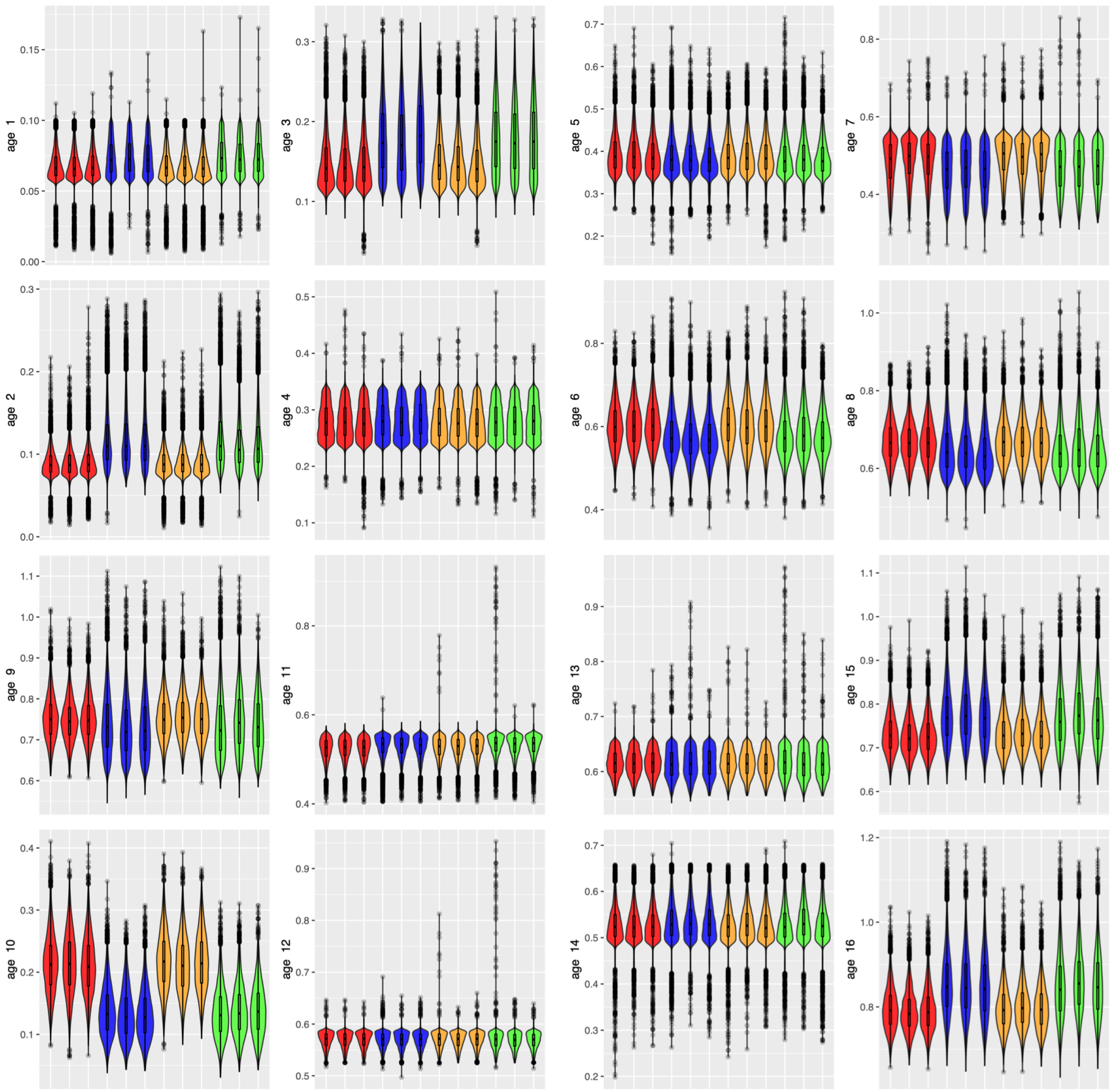


**Figure S3. Age of internal nodes 1-16 for 36 mammals from dos Reis et al.** (2012). Red and blue violin plots were obtained using the full phylogenetic likelihood (GTR+G4+I) under UGAM and WN rate priors respectively, while orange and green plots show the results for the two-step approach under, respectively, UGAM and WN rate priors. Computations were run in triplicates. Time is in units of million years ago.

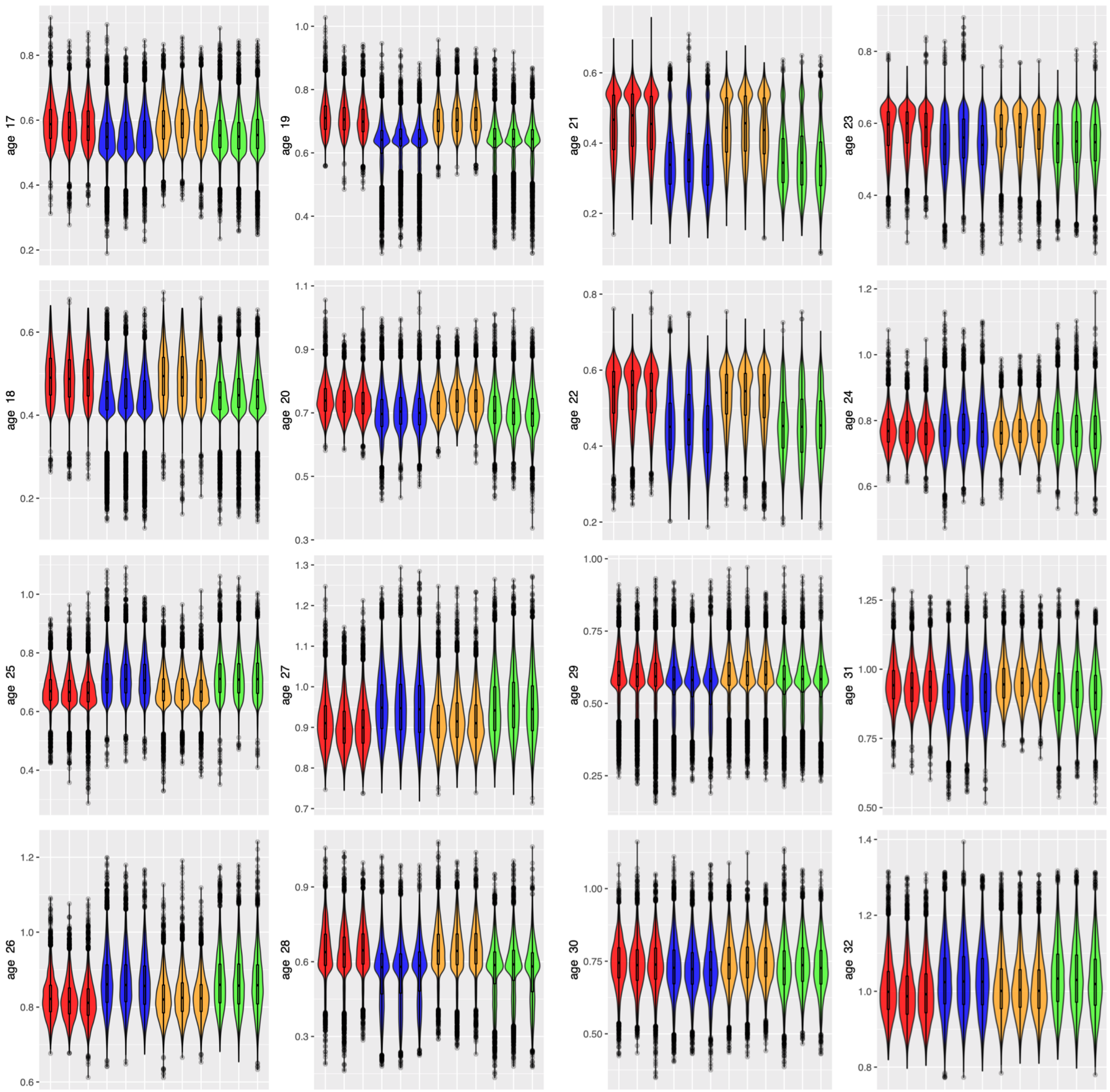


**Figure S4. Age of internal nodes 17-32 for 36 mammals from dos Reis et al.** (2012)**.** Red and blue violin plots were obtained using the full phylogenetic likelihood (GTR+G4+I) under UGAM and WN rate priors respectively, while orange and green plots show the results for the two-step approach under, respectively, UGAM and WN rate priors. Computations were run in triplicates. Time is in units of million years ago.

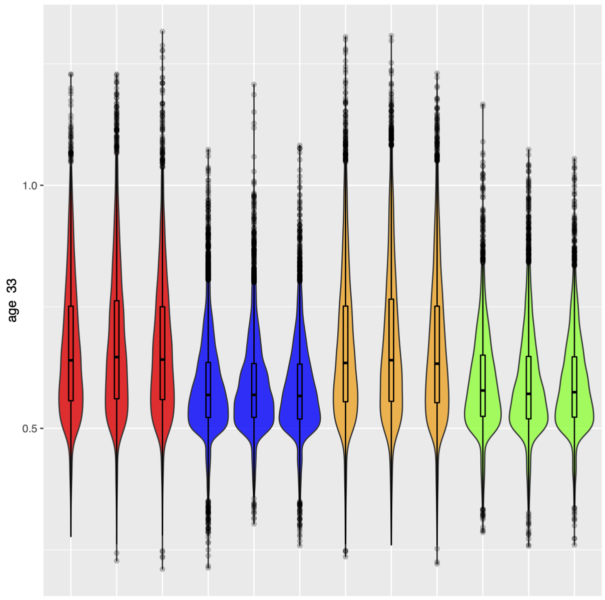


**Figure S5. Age of root node for 36 mammals from dos Reis et al. (2012).** Red and blue violin plots were obtained using the full phylogenetic likelihood (GTR+G4+I) under UGAM and WN rate priors respectively, while orange and green plots show the results for the two-step approach, respectively, UGAM and WN rate priors. Computations were run in triplicates. Time is in units of million years ago.

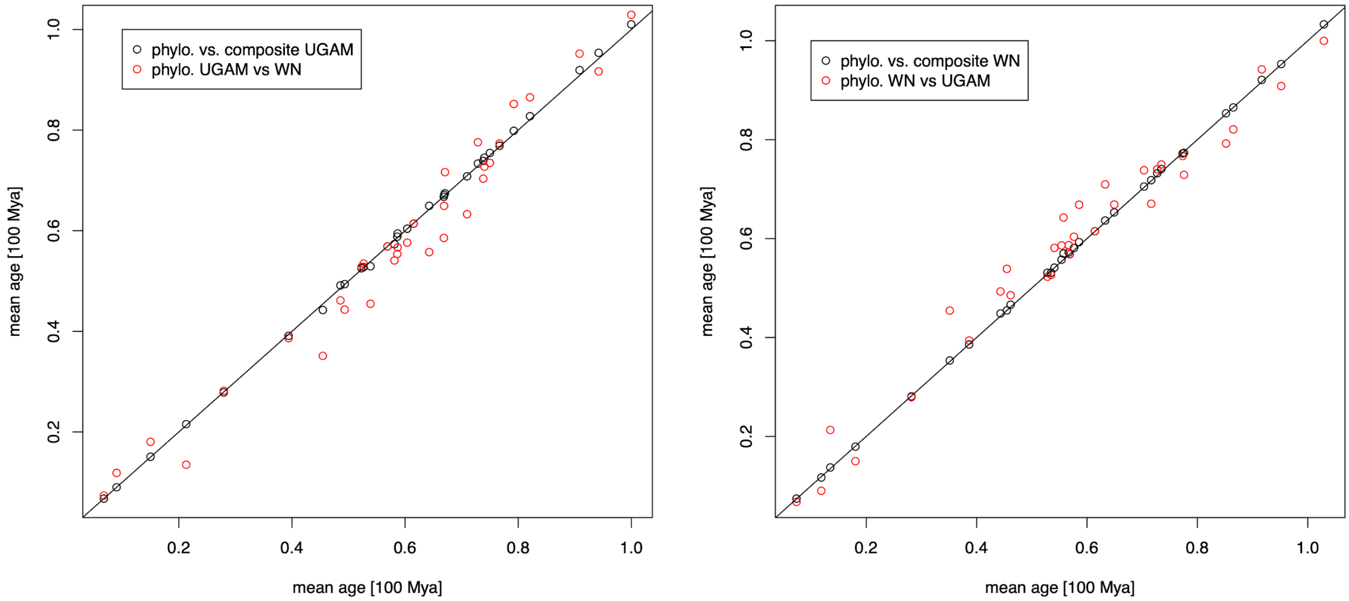


**Figure S6. Mean posterior age for internal nodes for 36 mammals from dos Reis et al.** (dos Reis et al. 2012)**.** Black circles reveal that the correlation between the full likelihood and the two-step approach is better than the correlation between different relaxed clock models, UGAM and WN, in red circles. The black line is the y=x line.

### On the informativeness of constraints

We have identified 3 factors that are expected to affect the informativeness of a constraint, and we discuss them here.

Firstly, whether a constraint is distal or proximal. In the general case, we expect proximal constraints to be more informative than distal constraints, because a proximal constraint implies several more distal constraints. In Fig. S7, we have noted inner nodes from after the root to the tips with letters A to I. If we have the proximal constraint that node D is older than node E, then it implies several additional more distal constraints: D older than H, but also B older than E, and B older than H. **For this reason, in general, a proximal constraint should be more informative than a distal constraint. In fact, its informativeness depends on the number of nodes ancestral to the older node of the constraint, and the number of nodes descendant to the younger node in the constraint.**

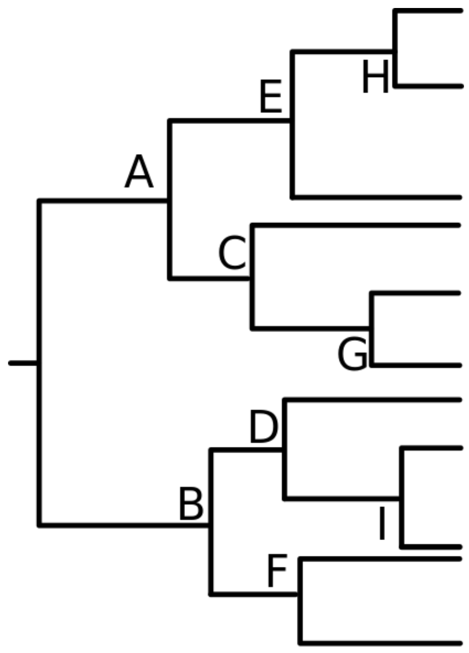


**Fig. S7: Schematic timetree to reflect on informativeness of constraints.** Inner nodes other than the root are annotated with letters A to I.

Secondly, the informativeness of a constraint depends on the underlying variation in the rate of molecular evolution. This rate variation makes dating a tree difficult. Therefore, constraints will be more informative if they apply to parts of a tree where rate variation is particularly severe, or behaves in a way that is not well fitted by usual models of rate variation.

Thirdly, the informativeness of a constraint depends on interactions with node age calibrations. Constraints can propagate the information provided by a node age calibration to another node. If there are few node age calibrations in a tree, or if these calibrations apply to nodes distant from the constrained nodes, then we expect that the constraint may be less informative than a constraint that involves a calibrated node.

To summarize, it would seem that the most informative constraints between two nodes should be such that:

- The older node has many ancestral nodes
- The younger node has many descendant nodes
- Rate variation in the vicinity of the constraint is pronounced
- A node in the vicinity of the constraint is calibrated, such that the constraint can propagate the information provided by the calibration

### Verification that the MCMC dating algorithm is well calibrated.

To check that the MCMC algorithm is well calibrated when the simulation and inference model are the same, we performed simulation and inference using RevBayes, using the same model for both steps. We simulated a tree topology with 102 tips under a birth-death process and simulated a sequence alignment of length 1000 sites under a Jukes-Cantor model of DNA sequence evolution and 20 categories of site rates according to a discretized Gamma distribution of shape and rate 0.3. Inference was performed under the same model, without relative node order constraints, with the same MC3 algorithm and the same moves as those used in our simulation analysis. We computed the Maximum A Posteriori tree and computed 95% High Posterior Density intervals on node ages. We find that they contain the true simulated node ages 96% of the time (97 nodes out of 101), as displayed on Supp. Fig. S8.

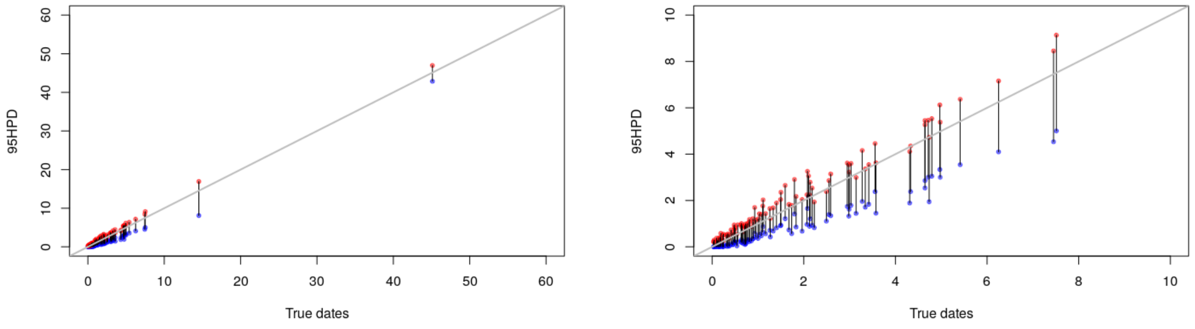


**Fig. S8: Estimation of node ages when the reconstruction model fits the simulation model.** Left: 95% High Posterior Density intervals are represented on the y axis for each true node age on the x axis. Red: upper 97.5% bound; blue: lower 2.5% bound. The line y=x is shown. Right: Zoom between node ages 0 and 10.

This result shows that the poor calibration observed on the simulations to test the effect of relative node order constraints is not due to deficiencies of the MC3 algorithm.

The simulation and inference scripts are available at https://github.com/Boussau/DatingWithConsAndCal/blob/master/Scripts/DatingRevScripts/simulateThenInferUGAM.Rev and https://github.com/Boussau/DatingWithConsAndCal/blob/master/Scripts/DatingRevScripts/endSimulateThenInferUGAM_InferUGAMParametersBactrianMC3.Rev.

### Timetrees for the Archaea for different maximum root ages

**Fig. S9 (following page):** We used the data including transfer derived relative age constraints from Davín et al. (2018) containing 62 species (Williams et al. 2017) with different maximum ages for the root (see main text). We highlight the two nodes on the archaeal tree discussed: i) the common ancestor of ammonia-oxidising (AOA, in green) Thaumarchaeota and the common ancestor of methanogenic Euryarchaeota (that is, the common ancestor Cluster 1 (red) and Cluster 2 (orange) methanogenic Euryarchaeota, corresponding to all Euryarchaeota except for the Thermococcus/Pyrococcus clade).

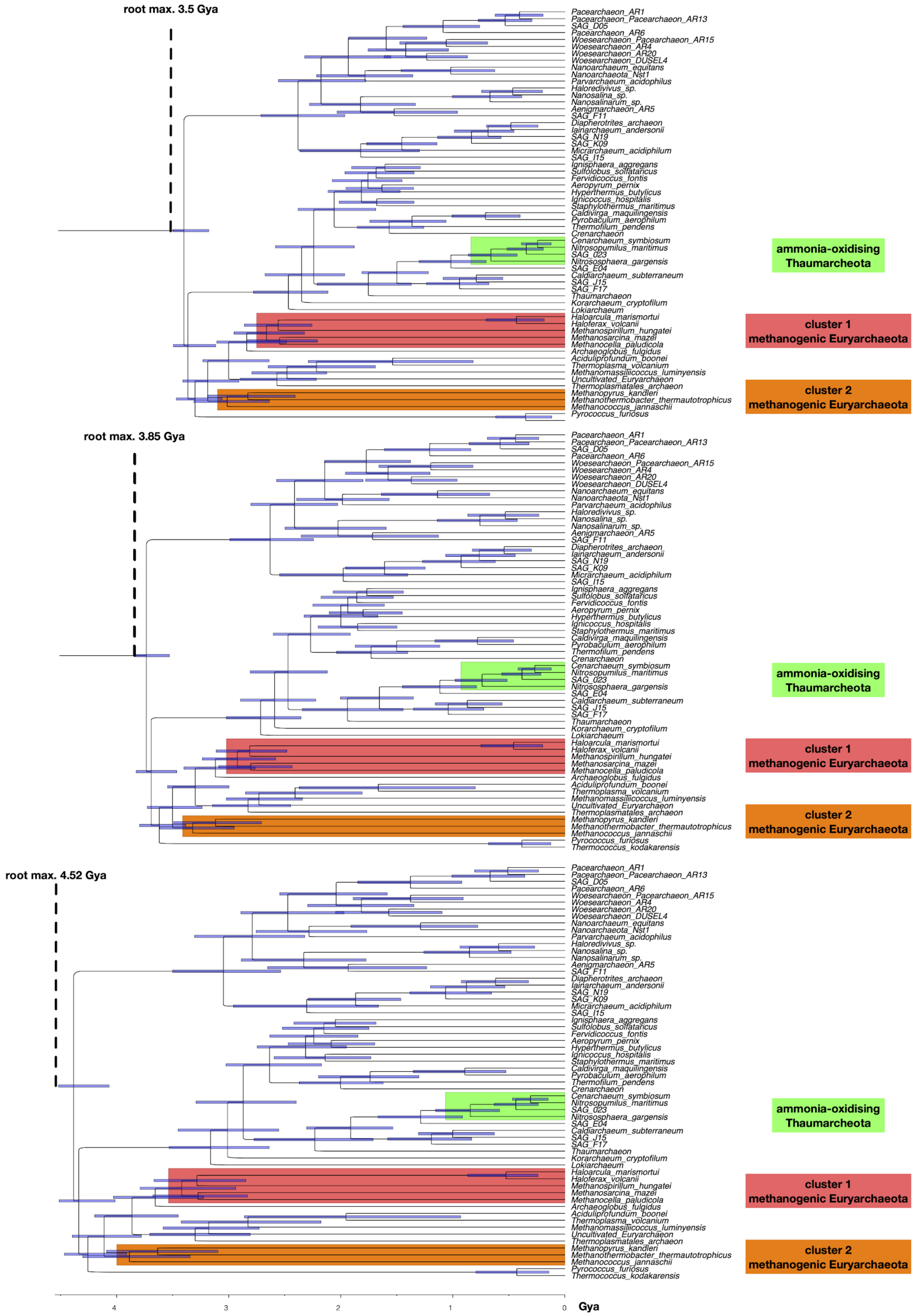
root max. 3.5 Gya
Pacearchaeon_AR1
Pacearchaeon_Pacearchaeon_AR13
SAG_D05
Pacearchaeon_AR6
Woesearchaeon_Pacearchaeon_AR15
Woesearchaeon_AR4
Woesearchaeon_AR20
Woesearchaeon_DUSEL4
Nanoarchaeum_equitans
Nanoarchaeota_Nst1
Parvarchaeum_acidophilus
Haloredivivus_sp.
Nanosalina_sp.
Nanosalinarum_sp.
Aenigmarchaeon_AR5
SAG_F11
Diapherotrites_archaeon
Iainarchaeum_andersonii
SAG_N19
SAG_K09
Micrarchaeum_acidiphilum
SAG_I15
Ignisphaera_aggregans
Sulfolobus_solfataricus
Fervidicoccus_fontis
Aeropyrum_pernix
Hyperthermus_butylicus
Ignicoccus_hospitalis
Staphylothermus_maritimus
Caldivirga_maquilingensis
Pyrobaculum_aerophilum
Thermofilum_pendens
Crenarchaeon
Cenarchaeum_symbiosum
Nitrosopumilus_maritimus
SAG_023
Nitrososphaera_gargensis
SAG_E04
Caldiarchaeum_subterraneum
SAG_J15
SAG_F17
Thaumarchaeon
Korarchaeum_cryptofilum
Lokiarchaeum
Haloarcula_marismortui
Haloferax_volcanii
Methanospirillum_hungatei
Methanosarcina_mazei
Methanocella_paludicola
Archaeoglobus_fulgidus
Aciduliprofundum_boonei
Thermoplasma_volcanium
Methanomassiliicoccus_luminyensis
Uncultivated_Euryarchaeon
Thermoplasmatales_archaeon
Methanopyrus_kandleri
Methanothermobacter_thermautotrophicus
Methanococcus_jannaschii
Pyrococcus_furiosus
ammonia-oxidising Thaumarcheota
cluster 1 methanogenic Euryarchaeota
cluster 2 methanogenic Euryarchaeota
root max. 3.85 Gya
Pacearchaeon_AR1
Pacearchaeon_Pacearchaeon_AR13
SAG_D05
Pacearchaeon_AR6
Woesearchaeon_Pacearchaeon_AR15
Woesearchaeon_AR4
Woesearchaeon_AR20
Woesearchaeon_DUSEL4
Nanoarchaeum_equitans
Nanoarchaeota_Nst1
Parvarchaeum_acidophilus
Haloredivivus_sp.
Nanosalina_sp.
Nanosalinarum_sp.
Aenigmarchaeon_AR5
SAG_F11
Diapherotrites_archaeon
Iainarchaeum_andersonii
SAG_N19
SAG_K09
Micrarchaeum_acidiphilum
SAG_I15
Ignisphaera_aggregans
Sulfolobus_solfataricus
Fervidicoccus_fontis
Aeropyrum_pernix
Hyperthermus_butylicus
Ignicoccus_hospitalis
Staphylothermus_maritimus
Caldivirga_maquilingensis
Pyrobaculum_aerophilum
Thermofilum_pendens
Crenarchaeon
Cenarchaeum_symbiosum
Nitrosopumilus_maritimus
SAG_023
Nitrososphaera_gargensis
SAG_E04
Caldiarchaeum_subterraneum
SAG_J15
SAG_F17
Thaumarchaeon
Korarchaeum_cryptofilum
Lokiarchaeum
Haloarcula_marismortui
Haloferax_volcanii
Methanospirillum_hungatei
Methanosarcina_mazei
Methanocella_paludicola
Archaeoglobus_fulgidus
Aciduliprofundum_boonei
Thermoplasma_volcanium
Methanomassiliicoccus_luminyensis
Uncultivated_Euryarchaeon
Thermoplasmatales_archaeon
Methanopyrus_kandleri
Methanothermobacter_thermautotrophicus
Methanococcus_jannaschii
Pyrococcus_furiosus
Thermococcus_kodakarensis
ammonia-oxidising Thaumarcheota
cluster 1 methanogenic Euryarchaeota
cluster 2 methanogenic Euryarchaeota
root max. 4.52 Gya
Pacearchaeon_AR1
Pacearchaeon_Pacearchaeon_AR13
SAG_D05
Pacearchaeon_AR6
Woesearchaeon_Pacearchaeon_AR15
Woesearchaeon_AR4
Woesearchaeon_AR20
Woesearchaeon_DUSEL4
Nanoarchaeum_equitans
Nanoarchaeota_Nst1
Parvarchaeum_acidophilus
Haloredivivus_sp.
Nanosalina_sp.
Nanosalinarum_sp.
Aenigmarchaeon_AR5
SAG_F11
Diapherotrites_archaeon
Iainarchaeum_andersonii
SAG_N19
SAG_K09
Micrarchaeum_acidiphilum
SAG_I15
Ignisphaera_aggregans
Sulfolobus_solfataricus
Fervidicoccus_fontis
Aeropyrum_pernix
Hyperthermus_butylicus
Ignicoccus_hospitalis
Staphylothermus_maritimus
Caldivirga_maquilingensis
Pyrobaculum_aerophilum
Thermofilum_pendens
Crenarchaeon
Cenarchaeum_symbiosum
Nitrosopumilus_maritimus
SAG_023
Nitrososphaera_gargensis
SAG_E04
Caldiarchaeum_subterraneum
SAG_J15
SAG_F17
Thaumarchaeon
Korarchaeum_cryptofilum
Lokiarchaeum
Haloarcula_marismortui
Haloferax_volcanii
Methanospirillum_hungatei
Methanosarcina_mazei
Methanocella_paludicola
Archaeoglobus_fulgidus
Aciduliprofundum_boonei
Thermoplasma_volcanium
Methanomassiliicoccus_luminyensis
Uncultivated_Euryarchaeon
Thermoplasmatales_archaeon
Methanopyrus_kandleri
Methanothermobacter_thermautotrophicus
Methanococcus_jannaschii
Pyrococcus_furiosus
Thermococcus_kodakarensis
ammonia-oxidising Thaumarcheota
cluster 1 methanogenic Euryarchaeota
cluster 2 methanogenic Euryarchaeota
4
3
2
1
0
Gya

Penel, Simon, Anne-Muriel Arigon, Jean-François Dufayard, Anne-Sophie Sertier, Vincent Daubin, Laurent Duret, Manolo Gouy, and Guy Perrière. 2009. “Databases of Homologous Gene Families for Comparative Genomics.” *BMC Bioinformatics* 10 Suppl 6 (June): S3.

Reis, Mario dos, Jun Inoue, Masami Hasegawa, Robert J. Asher, Philip C. J. Donoghue, and Ziheng Yang. 2012. “Phylogenomic Datasets Provide Both Precision and Accuracy in Estimating the Timescale of Placental Mammal Phylogeny.” *Proceedings. Biological Sciences / The Royal Society* 279 (1742): 3491–3500.

Williams, Tom A., Gergely J. Szöllősi, Anja Spang, Peter G. Foster, Sarah E. Heaps, Bastien Boussau, Thijs J. G. Ettema, and T. Martin Embley. 2017. “Integrative Modeling of Gene and Genome Evolution Roots the Archaeal Tree of Life.” *Proceedings of the National Academy of Sciences of the United States of America* 114 (23): E4602–11.
